# A META-ANALYSIS OF GUT MICROBIOME RESEARCH IN MALNOURISHED AFRICAN POPULATIONS: A NATURAL LANGUAGE PROCESSING APPROACH

**DOI:** 10.64898/2025.12.01.691114

**Authors:** Monica N Mweetwa, Paul Kelly, Joram M Posma

## Abstract

**Background:** Malnutrition still affects millions of children in Africa. Changes in the gut microbiome have been implicated in malnutrition, but there has been inconsistent nomenclature of microbes. This meta-analysis reviews the microbiome literature using natural language processing (NLP) methods.

**Methodology:** We searched PubMed for gut microbiome studies of undernourished children living in low-middle-income countries (LMICs). The primary analysis focused on continental coverage and study characteristics of microbiome research in sub-Saharan Africa. We also employed an NLP tool for normalising primary data from full-text publications in ss-Africa compared to other LMICs, and between diseased and healthy children.

**Results:** We identified 16 studies. Most studies were conducted in Malawi and characterised the faecal microbiome using 16S rRNA sequencing. For comparison, 18 studies conducted in Bangladesh, India, Pakistan and Peru were included. With this, we identified frequently reported microbes that were distinctly identified in sub-Saharan Africa and highlighted possible signatures of an undernourished faecal microbiome across the globe.

**Conclusion:** The consistent associations between elevated Pseudomonadota levels and severe acute malnutrition provides new insights into host-microbiome interactions in African contexts. However, the overlap between taxa associated with healthy and stunting underscores the need for further research to better inform potential targeted interventions in Africa.

## Section 1.01 Introduction

Malnutrition remains a critical and complex issue that plagues Africa (1). The continent’s diverse geographies, cultures (2,3), and socio-economic conditions contribute to the multifaceted nature of malnutrition (4,5). Despite notable progress in recent years, malnutrition persists, affecting millions of people, particularly children. Africa’s malnutrition problem is characterised by diverse phenotypes, including undernutrition, micronutrient deficiencies, and the growing prevalence of obesity (1). Undernutrition manifests in the form of stunting, wasting, oedematous malnutrition, and underweight among children, indicating a lack of proper dietary intake and access to essential nutrients (6). Micronutrient deficiencies are also prevalent due to limited access to diverse diets and specific geographically determined deficiencies such as selenium (1). Alongside undernutrition, there is a rising trend of obesity, particularly in urban areas, fuelled by factors like rapid urbanisation, changing dietary habits, and sedentary lifestyles (8).

There is evidence suggesting that the gut microbiome plays a significant role in the development and progression of malnutrition (9). The gut microbiome, a complex ecosystem of microorganisms residing in our intestines, influences nutrient absorption, immune function, and overall health (10). Understanding the composition and function of the microbiome has improved our knowledge of disease mechanisms by revealing how specific microbial communities and their metabolic activities influence host immune responses, inflammation, and metabolic pathways involved in conditions such as inflammatory bowel disease, metabolic disorders, and undernutrition (10–12). This detailed insight into which microbes are present and their roles has enabled the development and refinement of microbe-mediated therapies like faecal microbiota transplantation (FMT) (13–15), and microbiota directed feeds (16,17) aimed at restoring a healthy microbial balance in the gut.

Several studies have explored the correlation between malnutrition and the gut microbiome and excellent reviews on this subject are available (18–28). Despite this, the number of studies is still modest. There also is the difficulty of changes and inconsistencies in nomenclature (29–33) which make it difficult to systematically compare microbiome studies without expert knowledge. Advances in natural language processing (NLP) (34) techniques have facilitated the extraction of information from vast volumes of literature, and recently domain-specific tools have been developed for microbiome literature (35). However, to the best of our knowledge there has been no meta-analysis of African intestinal microbiome data focused on malnutrition besides Chibuye *et al*. (36) that focused on stunting. This meta-analysis aims to provide an (i) overview of gut microbiome studies conducted in undernourished populations from low-middle-income countries (LMICs) in sub-Saharan Africa (ss-Africa), and (ii) compare how frequently these microbes are reported in ss-Africa to other LMICs regions highlighting the application of natural language processing in literature mining to permit consistent identification of key taxa and improve summarisation of data.

## Section 1.02 Data and Methods

### (a) Search strategy

To build an overview of the gut microbiome landscape in malnourished populations in LMICs, a comprehensive search was carried out in PubMed for articles published up to 01/06/2024 as well as searching reference lists of reviews. The classification of LMICs was based on the World Bank classification at the time of the search. For the 2025 fiscal year, as well as in 2024 when the search was conducted, they used the income classification data from 2023 to classify LMIC countries (37). This search was restricted to PubMed because the NLP employed downstream could only be applied to files from PubMed Central (PMC). The terms of the search strategy are included in Supplementary data (*S1_Data*).

### (b) Eligibility criteria

Studies were eligible if they performed gut microbiota analysis and reported relative abundance of microbes at any taxonomic rank and were included if they characterised the gut microbiome in children or adolescents (<18yrs) with SAM defined by weight-for-length z-score (WLZ) < -3, middle upper arm circumference (MUAC) < 11.5cm, bilateral pitting oedema, stunting defined by length-for-age z-score (LAZ) < -2, wasting defined by weight-for-age z-score (WAZ) < -2 or children/adolescents (<18yrs) with environmental enteropathy (also called environmental enteric dysfunction) measured by a lactulose test, regardless of study design. The main outcome was microbial composition (at any timepoint if longitudinal sampling was done).

Abstracts of identified papers were reviewed for inclusion and studies were excluded if they (i) looked at obesity outside ss-Africa because they had no comparison group within ss-Africa (the group of primary interest), (ii) used a population that was African by descent, (iii) re-analysis of already published data (unless the secondary analysis included more microbial data from the same cohort, then the former was excluded), (iv) reported data from 1 undernourished child or characterised a single species, and/or (v) did not have a manuscript archived in PubMed Central (PMC). The PRISMA checklist for the meta-analysis is included (*S1 Table)*.

### (c) Data collection

The data collected included PMCID, DOI, title, publication year, country and region, aim of the study, setting of the study i.e. rural, peri-urban, urban etc., age of children included, sample size of the study, study design, category of undernutrition investigated, control groups included if any, sample type used to assay the gut microbiome, method of microbial characterisation and all microbes reported in the results sections of studies.

The selection of studies was done by 2 reviewers (MNM and PK). The extraction of microbiome data was carried out in duplicate using an NLP tool as detailed in the section below and verified manually by 2 reviewers (MNM and JMP) using TeamTat (38).

#### (i) Implementation of natural language processing (Qualitative synthesis)

Collection of all microbes reported in the results was automated using a named-entity recognition (NER) and named-entity normalisation (NEN) tool called Microbe Entity Linking Pipeline (MicrobELP) (39) that extracts the names of microbes reported in manuscripts and links these entities to taxonomic IDs as well as taxonomic rank and returns these in form of machine-readable JSON files. We used the published annotation pipeline (dictionary matching) with a precision over 99% as evaluated on a manually annotated test set of 288 articles (strict F1-score = 93.08, unpublished data, in submission). We adopted this approach to compare microbial composition reported. To do this, we converted HMTL files from PMC to BioC JSON files using Auto-CORPus (40). These JSON files were then used as input for the MicrobELP algorithm. The microbe search was restricted to the text in the results section of manuscripts to focus on the microbes that were most relevant to each study.

MicrobELP detects names and taxonomic identifiers that have bacteria, archaea or fungi in their lineage (super)kingdom to species. Here we manually verified all annotations and manually added taxa that had virus lineage. The NER algorithm was able to directly provide taxonomic identifiers (taxids) that were used as input for taxonomic classifications in Rv4.3 (41). This way, we were able to compare microbes reported in different studies using a consistent nomenclature. For example, Firmicutes and Bacillota, or Proteobacteria and Pseudomonadota, were recognised as the same taxon by the MicrobELP. Although the algorithm does identify ‘bacteria’ as being reported, it almost never refers to a finding e.g. ‘bacteria were significantly higher in the disease group’ but more generally talking about what was studied -e.g. bacteria opposed to archaea or fungi (42–49). A list of all microbes detected is included (*S2 Table*). Gehrig *et al*. (16) and Vonaesch *et al*. (49) reported both the gut microbiome of children and microbiome of animal models. These were manually screened to retain data in the results sections belonging to human participants only.

The taxids were used for phylogenetic trees construction of all microbes reported as well as comparative trees between regions (n = 34), health status (n = 11) and sequencing methodology (n = 34) using a custom pipeline available via GitHub (50). The packages used to do this were ‘ape’ (5.8) (51), ‘ggtree’ (3.12.0) (52), ‘taxonomizr’ (0.10.6) (53), and ‘ggplot2’ (3.5.1) (54).

#### (ii) Quantitative synthesis

The Fisher’s exact test was used in an exploratory analysis to determine if the frequency of microbes reported in at least 5 papers were significantly different by region. To account for potential biases, 1000 permutations were performed and permutated p-values < 0.05 were considered significant.

### (d) Assessment of bias and certainty

#### (i) Risk bias assessment

The studies included had multiple study designs so different Cochrane tools for risk of bias (ROB) assessment were used i.e. Cochrane’s risk of bias tool for randomized controlled and cluster randomized trials (RoB2) (55), Risk of Bias in Non-randomized Studies -of Exposures tool (ROBINS-E) (56) and the ROBIN-I tool (of Interventions) (57) were used for randomised control trials (RCTs) and cluster RCTs, observational studies and case control studies respectively. For case-control studies and non-RCTs, we considered age, antibiotic usage, breastfeeding, gender, and diet as potential confounding variables.

There were 12/34 studies with concerns stemming from bias in randomisation, missing data as majority of them were secondary analysis papers that did not include all participants from parent studies or had bias due to confounding variables not being accounted for. There were 7/34 studies with high ROB due to inappropriate analysis methods or bias from more than 1 key confounding variable. These were mainly observational studies which cannot completely be free of confounding bias. All details of this analysis for each study included (*S3 Table*).

Given the limited number of studies included in the meta-analysis, we opted not to exclude any studies based on the ROB assessment to ensure sufficient data for meaningful analysis. Moreover, the primary outcome focused on the reporting of microbes in manuscripts rather than effect estimates addressing the research questions of individual studies. For comparisons between cases and controls sub-analysis, data extraction was limited to non-randomised studies.

Characteristics of included studies in this meta-analysis is included (*S4 Table)* and graphical depictions of demographic features created using Python (58) are shown in the results and discussion.

## Section 1.03 Results and Discussion

The literature search yielded 575 articles from January 2013 to March 2024 in low-middle income countries across the globe. Thirty-four articles remained after manual screening for full texts and abstracts, of which 16 included populations from ss-Africa (*Figure 1*).

**Figure 1:**
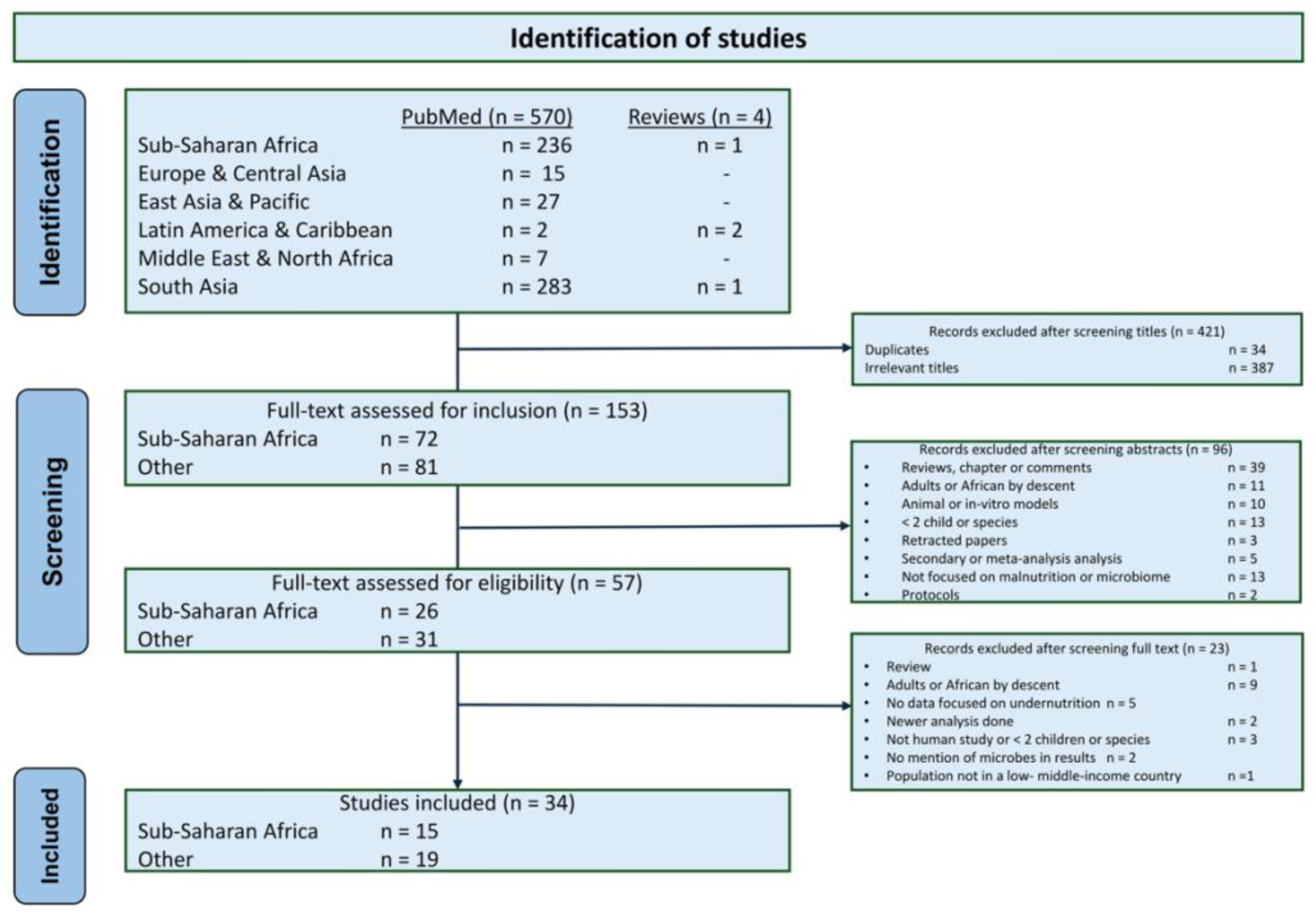
PRISMA exclusion diagram of studies surveyed in this meta-analysis.

## Section 1.04 Study characteristics of Sub-Saharan studies

Studies exploring the link between the gut microbiome and malnutrition in Africa have almost entirely been published in the last decade (*Figure 2A*). These studies have spanned the continent, but coverage has been very patchy (*Figure 2C*) and have focused on children with severe acute malnutrition (SAM) or stunting (*Figure 2D*). We did not find any studies looking at the microbiome with respect to micronutrient deficiencies or obesity in ss-Africa. These have been carried out in other LMIC regions such as Pakistan were they show that micronutrient supplementation in children under the age to 2 was associated with increased carriage of *Cryptosporidium* (59). With the increasing trend in childhood obesity (60), there is also need to expand the scope of malnutrition classes studied in ss-Africa, beyond undernutrition.

**Figure 2:**
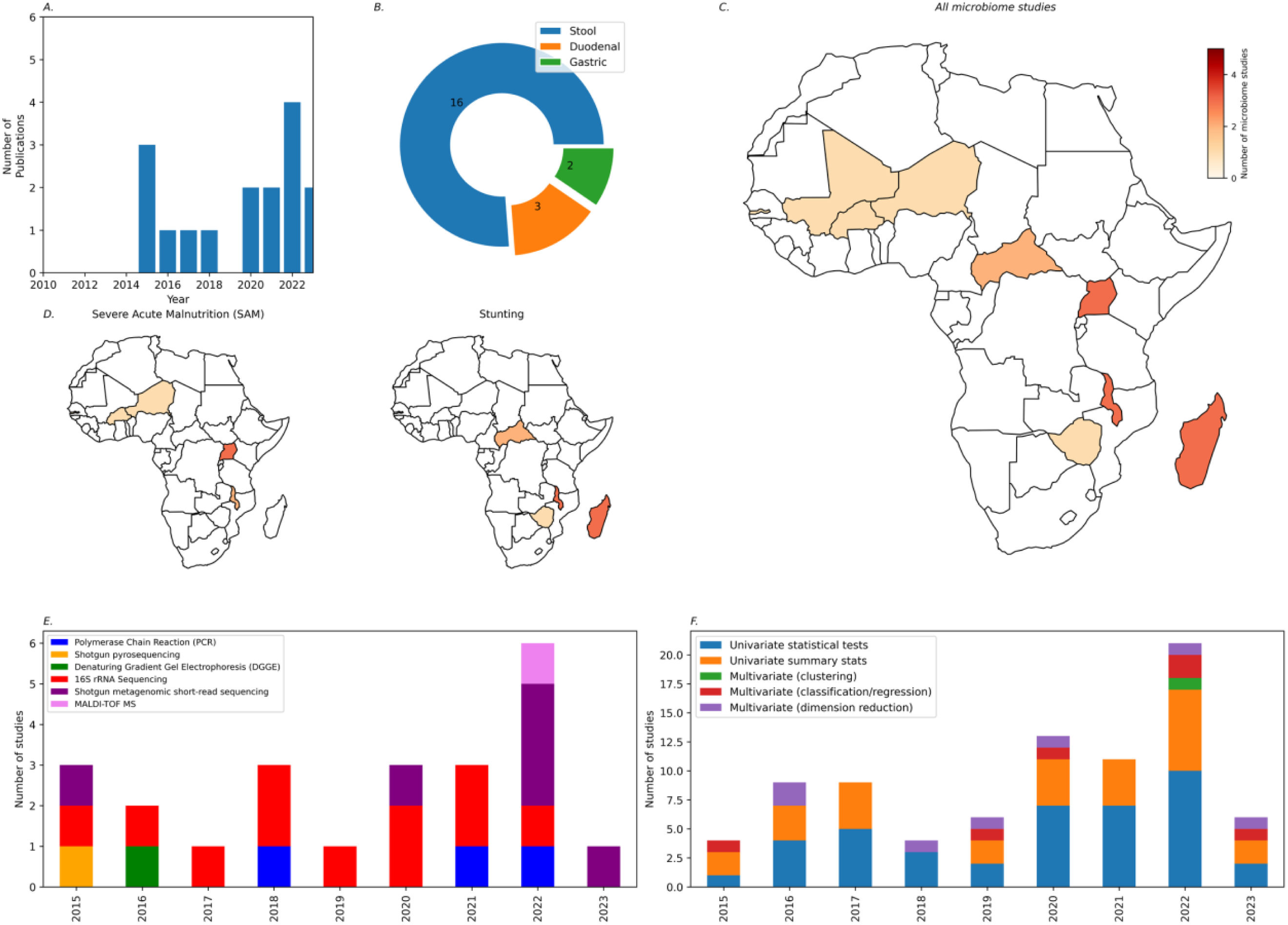
Overview of gut microbiome studies carried out in malnourished children in LMICS in Africa. (A) Number of publications per year. (B) Type of sample used as proxy for gut microbiome. (C) Geographic distribution of malnutrition studies looking at the gut microbiome. (D) Geographic distribution by type of malnutrition studies conducted with African participants (SAM or stunting). (E) Type of assay used to measure the microbiome. (F) Statistical analysis methods used *(some studies used multiple assays to detect the microbiome and statistical methods for analysis)*.

### (a) Sample types

Faecal samples have been the primary source for studying the gastrointestinal (GI) microbiome used in all the of studies reported due to ease of collection (*Figure 2B*). While they provide valuable insights into the overall composition of gut bacteria, they do not fully represent the microbial communities present along the entire GI tract (61). There were 3 studies looking at the small intestinal regions of the gut however, these studies were all carried out in the same population (Central Africa republic and Madagascar), making it difficult to extrapolate these findings to other regions. No studies analysed intestinal mucus or mucosal biopsies. These tissue samples directly reflect the microbial communities in close contact with the gut epithelium, which is critical for understanding host-microbe interactions in malnutrition where malabsorption and epithelial abnormalities are primary characteristics (62). Biopsies can also help identify local variations in the microbial composition, which faecal samples cannot capture accurately. Collection of such samples can be challenging especially in vulnerable children, however newer, less invasive technologies of sampling the proximal gut (63–65) are on the rise therefore more coverage of these sample in the near future is anticipated.

The oral cavity is the first point of contact for many microbes that enter the GI tract, and its microbiota has been well studied. Known members of the oral cavity, such as *Streptococcus salivarius*, *Haemophilus* spp., *Neisseria* spp. and *Veillonella* spp., have been reported in small intestine and stool of stunted children (48) and have been shown to cause lipid malabsorption in mouse models (49). It has been suggested that decompartmentalization of the microbiome, with oral taxa aberrantly present in the lower parts of the gut could contribute towards manifestation of stunting; however, the characterisation of the oral microbiome within the oral cavity in malnourished individuals is relatively underexplored. There is also a dearth of data on the healthy microbiome of the stomach and proximal small intestine, and how impairments of the gastric acid barrier might permit decompartmentalization.

### (b) Sequencing technologies

The emergence of Next-Generation Sequencing (NGS) technologies such as Illumina sequencing, allowed for high-throughput analysis. In ss-Africa, these NGS technologies have enabled researchers to conduct more extensive and detailed studies of the gut microbiome in malnutrition across various populations and regions as well as study possible functional pathways associated with malnutrition which were previously not known with PCR technologies. More recent advances have introduced long-read sequencing technologies, such as Pacific Biosciences (66) and Oxford Nanopore Technologies (67). These methods offer longer read lengths, which facilitate the assembly of complex microbial genomes, including repetitive regions, with high throughput. In the context of malnutrition in ss-Africa, long-read sequencing has not been utilised but provides new possibilities for studying less well-characterised and potentially novel gut microbial species that may play a role in malnutrition outcomes.

PCR based methods enable the identification specific microbial genes, notably sequencing of the 16S rRNA region has dominated gut microbiome research (*Figure 2E*) despite not covering archaea organisms or viruses which are also present in the gut. As sequencing technologies have continued to evolve, malnutrition studies in Africa have increasingly adopted integrative approaches, combining metabolomics data. A multi-omic approach to characterising the gut microbiome provides a more comprehensive understanding of the functional capabilities and interactions within the microbial community as well as interactions with the surrounding milieu (68,69).

### (c) Analytical tools implemented

In the early stages of malnutrition studies in Africa, researchers mainly relied on exploratory statistical methods to investigate the role of the gut microbiome (*Figure 2F*). Descriptive statistics, such as mean, median, standard deviation, including ecological measures such as alpha and beta diversity were used to understand the overall microbial composition in malnourished populations compared to healthy controls (45,70,71). These preliminary studies laid the groundwork for identifying potential microbial biomarkers associated with malnutrition. As the field progressed, researchers adopted univariate statistical methods to determine differential abundance of microbial taxa between malnourished and healthy individuals. Techniques like Student’s t-test and analysis of variance (ANOVA) were utilised to identify specific taxa that showed significant differences in abundance between the two groups. However, these approaches are not ideal because microbiome data does not usually meet the method assumptions of homogeneity of variance and normality which are key assumptions for these analyses (68).

To overcome the limitations of univariate methods, multivariate techniques such as Principal Component Analysis (PCA) and clustering algorithms are increasingly being adopted. PCA enables researchers to reduce the dimensionality of sequencing data while capturing the most significant variations between malnutrition states (72). Clustering algorithms, such as hierarchical clustering and k-means clustering, allowed the identification of microbial communities associated with specific malnutrition phenotypes, aiding in the classification of individuals into distinct subgroups (43,73). More recently, algorithms like Random Forest have been used to classify malnutrition outcomes based on complex microbial interactions (12,73). Some studies also use a combination of methods that create an advantage of teasing out important features that would otherwise be missed if only one method was employed [29], [33], [36], [63], [64], [65].

## Section 1.05 Leveraging natural language Processing (NLP) to understand microbiome literature

We employed an NLP tool called MicrobeELP to extract microbial data from the text of results sections of the literature. The entire annotation process took under two hours to complete and provided taxonomic IDs, which were then used to visually assess microbial patterns. This was significantly faster than manual annotation, which took over a week, highlighting how MicrobELP greatly enhanced the efficiency of the meta-analysis. One limitation of using MicrobELP is the lack of viral annotations which were added manually (independently by two researchers). TeamTat was used in collaborative mode to resolve any disagreements (38).

MicrobELP generated 238 correctly identified unique taxa and 6 ambiguous annotations. Ambiguous annotations included Actinobacteria, Bacteroidetes and Spirochaetes which could either be classified under the Phylum or Class categories and *Bifidobacterium breve* (*B. breve*) which when abbreviated could have multiple matching taxonomic ids; these were correctly annotated during manual verification. The manual verification of annotations also added 69 taxa from several virus and eukaryotic superkingdoms, and bacteria that were wrongly spelt in papers which were not recognised by MicrobELP.

In total, 308 unique taxa were identified in human samples across 34 studies included of which 198 were detected in ss-African studies (S2 Table). The other regions included in this analysis for comparison purposes were South Asia and Latin America & Caribbean.

Approximately 39% of these microbes were reported in at least two studies (S2 Table) regardless of region. The most prevalent taxa, Pseudomonadota (Class) and *Prevotella* (Genus) were reported in 59% (20/34) of all the studies. The top taxa reported in at least 20% of all studies were Pseudomonadota (Phylum, 20/34), *Prevotella* (Genus, 20/34), Bacillota (Phylum, 19/34), *Bifidobacterium* (Genus, 16/34), Bacteroidota (phylum, 16/34), Actinomycetota (Phylum, 16/34), *Streptococcus* (Genus, 14/34), *Escherichia coli* (Species, 14/34), *Lachnospiraceae* (Family, 11/34), *Clostridium* (Genus, 10/34), *Shigella* (Genus, 10/34), *Lactobacillus* (Genus, 10/34), *Enterobacteriaceae* (Family, 9/34), *Bacteroides* (Genus, 9/34), *Escherichia* (Genus, 9/34), Oscillospiraceae (Family, 8/34), *Megasphaera* (Genus, 8/34), *Bifidobacterium longum* (Species, 8/34), *Campylobacter* (Genus, 7/34), *Veillonella* (Genus, 7/34), *Klebsiella* (Genus, 7/34), *Blautia* (Genus, 7/34) and *Faecalibacterium* (Genus, 7/34) (*Figure 3A*).

**Figure 3:**
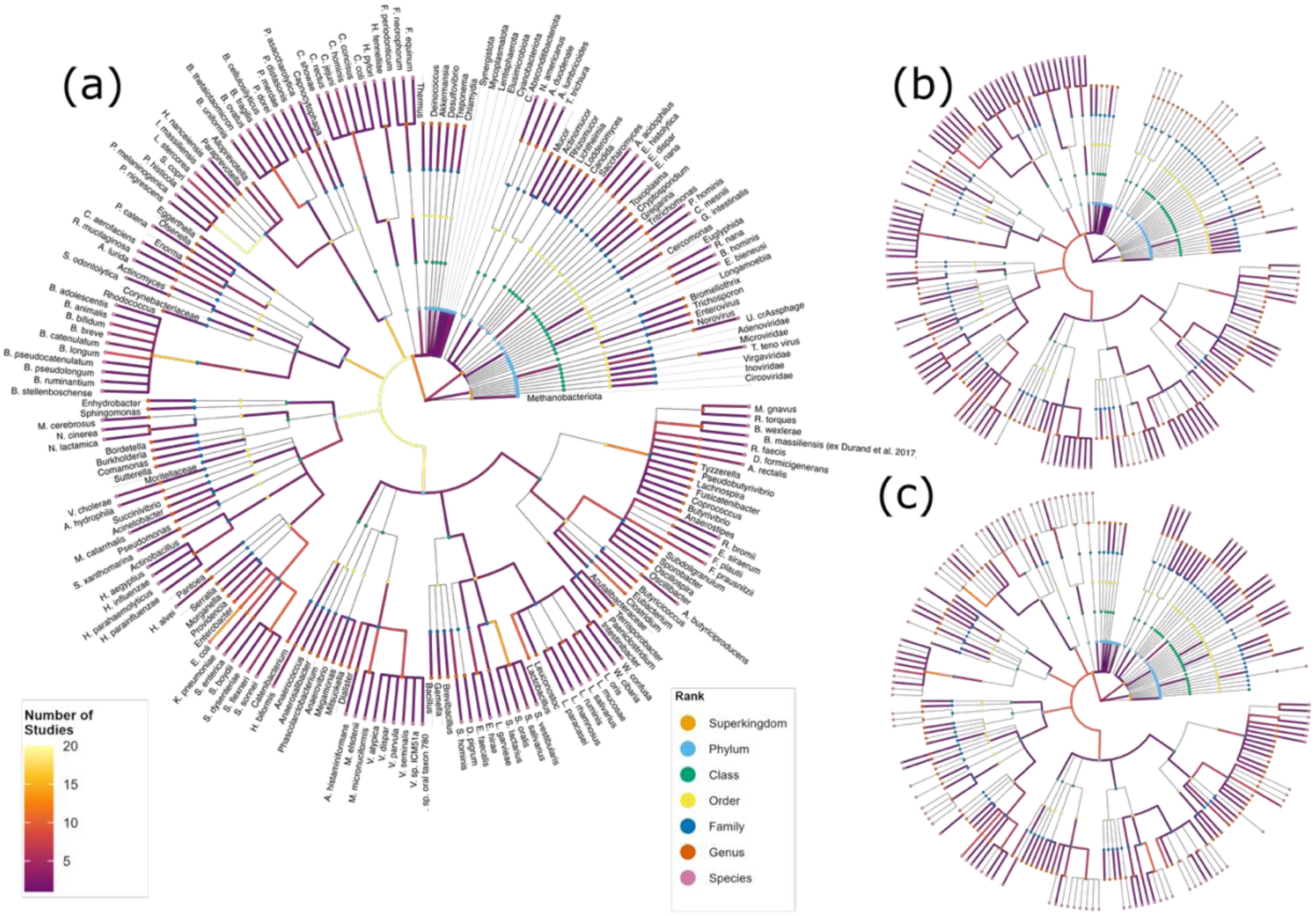
Phylogenetic diversity of microbes found in malnutrition studies. (A) Phylogenetic ranking of microbes reported in studies **across the globe** and prevalence of each microbe in the tree. Each node represents a microbe, coloured by rank. The colour of edges connected to a node in the tree represents the number of studies reporting it. (B) Phylogenetic ranking and prevalence of microbes reported in sub-Saharan Africa. (C) Phylogenetic ranking and prevalence of microbes reported in LMIC’s outside sub-Saharan Africa i.e. Asia and South America. The colour scales for (B) and (C) are the same as (A).

### (a) Are the associations with undernutrition unique within sub-Saharan Africa?

We used Fisher’s exact test to determine if there was a significant difference in reporting of microbes within ss-Africa compared to other countries and noted that *Bifidobacterium* (p = 0.04) and *Faecalibacterium* (p = 0.01) were reported at significantly lower frequency within ss-Africa compared to other regions. *Bifidobacterium* has been linked to breastfeeding (78,79). Within ss-Africa, use of amoxicillin for treatment of SAM has been shown to decrease the proportional abundance of *Bifidobacterium* (80) and has been linked to elevated levels of alpha-1 antitrypsin (AAT) in stunted children (49), similar to findings in Bangladesh (81). In South Asia, an increased the proportional abundance of *Bifidobacterium* spp. such as *B. longum* and *B. breve* has been linked to increasing age (82) as well as improved weight-for-age (WAZ) (83) and nutritional status (84). In all studies included, there was no analysis of breastfeeding status and microbial composition. Barratt *et al*. (85) showed that supplementation with *B. infantis* (a microbe highly associated with breastmilk) improved weight gain and reduced inflammation in children with SAM. There is a need to explore the relationship between breastfeeding and malnutrition in ss-Africa.

The *Faecalibacterium* genus was not reported in any of the ss-African countries though species such as *Faecalibacterium prausnitzii* have been reported. In other regions such as Asia, this genus has shown both positive and no association with nutritional index in Indian children (84,86). In other studies, this is simply amongst the most prevalent taxa reported (81–83). Other microbes demonstrated consistent detection frequencies across regions.

Bacillota (formerly Firmicutes) was the most frequently reported phylum. This phylum exhibits a wide metabolic repertoire, including fermentative metabolism and production of bioactive compounds (87). Many Bacillota species, such as *Bacillus*, can form spores, which are highly resistant to environmental stressors. This makes them more likely to persist and be detected in environmental and host microbiomes (88,89). In these studies, the Bacillota phylum was most abundantl reported phylum in stool samples cross all regions. Members of this phylum such as Lachnospiraceae families had variable associations with malnutrition by region. Lachnospiraceae has been associated with transition from SAM to healthy nutritional status in Gambian children (90) and was elevated post discharge in children treated for SAM with a probiotic (*Lactobacillus rhamnosus* and *Bifidobacterium animalis*) (91) in Uganda. However, reductions in Lachnospiraceae proportional abundance have been reported after dietary interventions with resistant starch (46) and rice-bran supplementation (92). Outside Africa, this family has been reported at higher proportional abundances in wasted and stunted children compared to seemingly healthy counterparts in Asian populations (83,93,94). This regional variability suggests that the role of Lachnospiraceae in malnutrition may depend on local dietary patterns, environmental factors, and co-existing microbial communities.

In ss-Africa, Pseudomonadota was amongst the most abundant phyla reported (42,44,46,47,90,95–98) and was significantly associated with infant mortality (42) in children with SAM. Pseudomonadota did not seem to be alleviated by supplementation with resistant starch (46). Most known pathogens such as *Salmonella* spp.*, E. coli, Shigella* spp. *Vibrio cholerae* and *Neisseria* spp. are members of this phylum so this hints at the likelihood of high pathogen burden in children with malnutrition. Pseudomonadota is also highly abundant in children with SAM in Bangladesh (99) and adults with metabolic disorders such as IBD (100) mostly caused by members of the *Enterobacteriaceae* family. Interestingly, Pseudomonadota was inversely associated with environmental enteropathy (EE) severity in stool samples (47) which was driven by a reduced abundance of *Klebsiella* and *Succinivibrio* but dominated the duodenal bacterial profile of stunted children (48,49) from a similar geographical location driven by higher prevalence of *Shigella*, *Campylobacter*. EE is strongly linked to stunting so this might further emphasise the difference between the stool microbiome and upper GI microbiome or suggest that the GI microbiome in EE is distinct from stunting. These variations in the abundance of Pseudomonadota in different cohorts likely reflects the need to look deeper than the phylum level for signatures that may have a causative relationship with SAM.

The Actinomycetota phylum, another dominant group in undernourished children, showed variable associations. It decreased with age in both undernourished and healthy children across Pakistan, India, and Peru (82,82,101), while some studies found it enriched in healthy children (84). The associations in both instances were driven by *Bifidobacterium*.

The *Prevotella* genus was found to be ubiquitous in African malnutrition and was neither significantly different to controls (46) nor related to severity of malnutrition (97). Individual *Prevotella* species detected were each reported in a single study so we cannot identify common features within malnutrition sub-types. The associations of *Prevotella* are complex and context-dependent, as its role can vary based on several factors including disease type and geographical region. Increased levels of *Prevotella* have been associated with conditions like inflammatory bowel disease (IBD) (102). Similarly *Prevotella* has been associated with stunting in Indian children (103). On the contrary, *Prevotella* spp. have been associated with recovery from moderate acute malnutrition in Pakistani children assisted by microbiota-directed feeds (17). Also, *Segatella copri*, commonly classified as *Prevotella copri* (*P. copri*) has been linked to improved glucose metabolism and lower risk of metabolic disorders such as type-2 diabetes in China (104). In Africa, however, it was highly abundant in two studies looking at stunted children (48,105) and Nigerians with type-2 diabetes (106). *P. histicola, P. melaninogenica* and *P. nigrescens* have also been reported in children with stunting in central and southern Africa (48,105). *P. melaninogenica* is considered a commensal of the oral mucosa in the early stages of life (107) whereas *P. histicola* strains have been shown to have anti-inflammatory properties in animal models of multiple sclerosis and rheumatoid arthritis (108,109). The detection of *Prevotella* in ss-Africa could also be attributed to the high fibre diet (110,111), (112) common to most regions of Africa. The complexity of its various associations with disease and health indicates that further typing at the sub-species level may be required.

*E. coli* was not unique to any region and was only reported in stool samples. *E. coli* was very prevalent regardless if they were healthy or malnourished (43,95,113), further suggesting that high pathogen carriage in communities with malnourished children. In SAM particularly, an increased proportional abundance of *E. coli* has been correlated to increasing severity of SAM (90).

*E. coli* can further be classified based on presence of pathogenicity islands, virulence factors and location of disease manifestation (114). Some subtypes of *E. coli* can also be commensal in the gut. 16S rRNA gene sequencing does not capture the regions in the *E. coli* genome that determine this difference which could possibly undermine the impact of *E. coli* in malnutrition. Pathotypes are also often omitted from whole genome sequencing data as well. Long read sequencing might be able to mitigate this but has so far not been applied. Besides this, targeted PCR designed to recognise the pathotype specific regions can also be used. The Afribiota group (43) used qPCR to show that the aaiC gene, a virulence gene of enteroaggregativ*e E. coli (EAEC)* was negatively correlated with height-for-age (HAZ) z-score in both stunted and healthy children in Madagascar. We have previously (115) shown using a Luminex platform that Zambian children with enteropathy had high enterotoxigenic *E. coli* (ETEC) regardless of malnutrition status. Both EAEC and ETEC has been associated with marker of enteropathy such as calprotectin and Reg1B in children from Bangladesh (116). In Pakistan, Aziz *et al*. (93) showed that microbial diversity can vary when *E. coli* is grouped by pathotype in Pakistani children that are stunted and the most prevalent pathotype detected was EPEC. There is still work to be done to understand the pathways through which *E. coli* pathotypes contribution towards malnutrition and if these are region specific.

We also noted that studies report depletion of *Lactobacillus* and elevation of *Veillonella* were in undernourished populations in ss-Africa. *Lactobacillus* is has been positively associated with birth weight and is dominant in the vaginal microbiome of pregnant mums (113,117) whereas *Veillonella* species have consistently been linked to undernutrition in Asian populations (22). *Veillonella* abundance is also associated with childhood obesity in western settings (118–120) but this was not seen in African cohorts because there are no published studies of obesity in childhood in Africa.

### (b) Beyond bacteria

Research on the gut microbiome in undernourished populations has predominantly focused on bacteria, with limited attention to viruses, archaea, and protozoa. However, the few studies exploring these non-bacterial microbes have revealed compelling insights.

For instance, Popovic *et al*. (59) demonstrated that micronutrient supplementation can modulate the eukaryome in children with inadequate growth in Pakistan. They observed increases in fungal genera such as *Rhizomucor*, *Actinomucor*, and *Mucor*, which were normalised with the addition of zinc to the supplement. Also, Reyes *et al*. (73) found that viral signatures in twins discordant for SAM differed significantly from those in concordant twin pairs, suggesting that viral dynamics may play a critical role in malnutrition outcomes.

These findings underscore the need for further investigation into the roles of non-bacterial microbes in undernutrition, as they may hold key insights into microbial interactions and gut health.

### (c) Is there a dysbiosis in malnutrition?

The analysis above summarised all studies regardless of study design i.e. some associations seen were a result of analysis on a continuous scale of malnutrition, e.g. length-for-age z-score (LAZ) or WAZ. To have a clearer picture of what has been observed between malnourished and non-malnourished children, we analysed the microbes reported in 11 studies that had a healthy control group irrespective of region. These were mainly in stunted children (Table S3) revealing notable trends. *Allisonella histaminiformans (Species)*, *Bifidobacterium (Genus)*, *Bifidobacterium longum (Species)*, *Blautia (Genus)*, Clostridiaceae (Family), *Collinsella (Genus)*, *Faecalibacterium (Genus)*, *Fusicatenibacter (Genus)*, *Intestinibacter (Genus)*, *Megamonas (Genus)*, *Roseburia (Genus)*, and Synergistota (Phylum) were associated with healthy status in at least 2 studies (Figure 4A). Species within these groups are short-chain fatty acids (SCFAs) producers strengthen the gut epithelial barrier, provide energy to colonocytes, and reduce intestinal inflammation (121–125).

**Figure 4:**
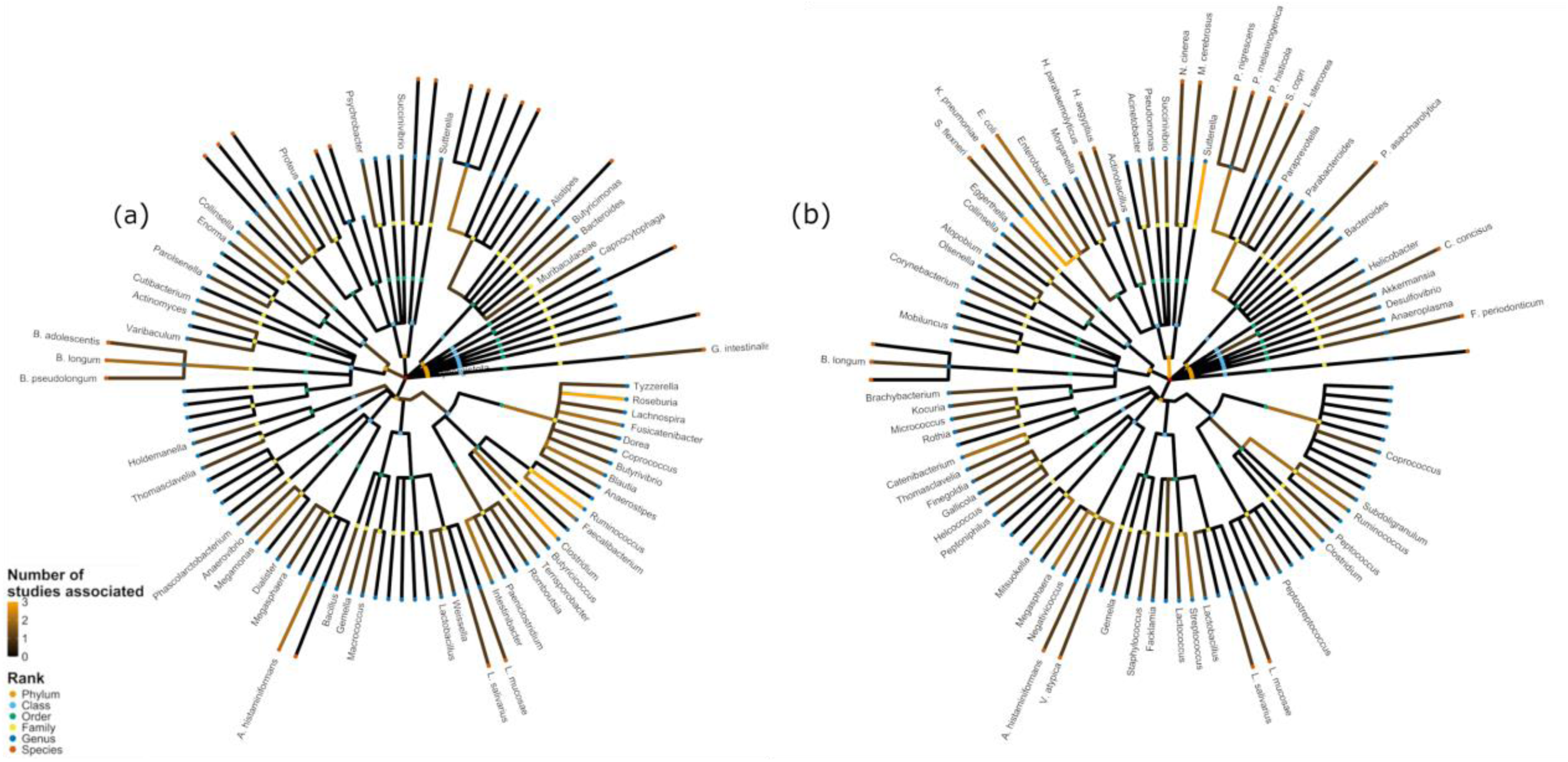
Phylogenetic trees showing microbes found associated with either undernutrition in (a) or (b) healthy status in LMIC populations.

At least 2 studies report associations between undernutrition and *Catenibacterium (Genus)*, *E. coli (Species)*, *Megasphaera (Genus)*, *Mitsuokella (Genus)*, *Oscillospiraceae* (Family), *Porphyromonas (Genus)*, *Prevotellaceae* (Family), Pseudomonadota (Phylum), *Shigella (Genus)*, *Streptococcus (Genus)*, *Sutterella* (Genus) and *Veillonella (Genus)* (Figure 4B). This suggests that there might be microbes whose increased proportional abundance promotes undernutrition. Pathogens such as *E. coli* (see above) and *Shigella* (genetically indistinguishable from E. coli) are known to cause intestinal infections. Most of these microbes are detected at family or genus level so functional prediction can be challenging. Deeper metagenomic research focused on these microbes might provide better insight to the potential roles of these microbes in promoting undernutrition.

Certain taxa were associated with both healthy and undernutrition statuses in at least two studies. These include *Clostridium*, *Escherichia*, Lachnospiraceae, *Prevotella* and *Ruminococcus*. Such dual associations may reflect context-dependent roles.

### (d) The influence of methodology on microbes reported

We compared the taxa reported by different sequencing methods classified as targeted sequencing (PCR, 16S, or 18S) (n = 24) and whole genome sequencing (WGS) (n = 6) (*S1Figure*). Papers reporting WGS-based data included viruses but not archaea while targeted approaches did not report any viruses except for *Norovirus* which is likely due to methodological focus. Additionally, WGS studies mainly reported taxa at species, genus, and phylum levels, while targeted sequencing studies reported across a broader range of taxonomic ranks.

Within targeted 16S rRNA sequencing studies, the choice of hypervariable region targeted also contributes to variation in reported microbes. Among the 16S-only studies (n=16), 13 targeted the V3-V4, V4, or V4-V5 regions, while 3 targeted V1-V2 or V1-V3 regions.

## Section 1.06 Conclusions and Future Perspectives

There has been steady growth of microbiome research in malnourished African populations. However, the revision of microbial taxonomy e.g. the 2021 revisions (126) results in inconsistent naming of microbes reported making it difficult to compare results between studies. Natural Language Processing (NLP) are methods used to identify, extract, and transform large collections of texts (34) and offer a way to speed up the analysis process and enhance accuracy. We employed MicrobELP, a domain-specific bio-NLP tool that can extract the names of microbes reported in manuscripts and map them to taxids in the NCBI database with > 98% precision. This automation significantly enhanced the comparability of findings across studies with varying nomenclature, thus streamlining data synthesis and interpretation. One notable aspect to emphasise is the need to expand tool coverage to process documents beyond those accessible in PMC via Auto-CORPus (40) and inclusion of microbes in the virus super kingdom in MicrobELP’s dictionary for improved coverage. Another limitation to the interpretation of the current NLP output is the lack of distinction between sequencing methods (no method exists that recognises sequencing technology). For example, the low counts of viruses reported may not be due to absence of these taxa in the microbiome, but a limitation of detection methods used to quantify each component of the microbiome. Methodological biases can be introduced at multiple stages, from sample collection and DNA extraction to sequencing and data analysis. Despite efforts to standardize protocols-such as the development of guidelines like STORMS (127) and best practice recommendations (128), bias remains a persistent issue that can distort observed microbial community composition. Our study acknowledges these limitations and contributes to this evolving field by highlighting how different methodological choices influence microbiome outcomes. Overall, integrating NLP into our analysis increased the efficiency of the review process, allowing for a more focused analysis of paper content and narrowing down features in the paper to focus on. With improvement, this NLP tool has the potential to improve the review process of microbiome literature in a systematic manner as it cross-references the text to known identifiers. Unlike large language models it does not generate text and is highly precise; however, its recall can be improved by means of using it to train a deep learning model that may be able to generalise more effectively.

We focused on African studies as there has been no summarisation of African malnutrition-microbiome literature in a systematic manner. The comprehensive body of evidence presented in this paper show that the gut microbiome is affected by changes in nutritional status and gives an indication to what microbes are likely to arise because of this. Overall, elevated levels of Pseudomonadota were consistently linked to malnutrition in Africa, especially SAM. EE is often associated with stunting and our analysis showed that a stunted gut microbiome in very similar to an EE gut microbiome. The most prevalent microbes associated with malnutrition were similar across LMICs around the globe apart from *Feacalibacterium* and *Bifidobacterium*, which vary geographically The findings from the healthy vs malnutrition analysis highlighted a spectrum of microbial associations with health and undernutrition which could serve as potential microbial biomarkers to distinguish health and undernourished children. However, the overlap in taxa associated with both states suggests that microbial signatures are context-dependent rather than universally fixed, indicating that while the microbiome holds promise for clinical diagnostics, its utility may vary across populations or settings.

While published research provides valuable insights, it also underscores the necessity for further investigations to unravel the complex interplay between host-microbiome interactions and malnutrition, particularly in neglected areas of the African context. These areas include understanding the role of the gut virome and mycobiome, as well as exploring gut microbial relationships with micronutrient deficiencies and childhood obesity. Such endeavours hold promise for advancing our understanding of malnutrition and informing targeted interventions to improve health outcomes in affected populations.

## Section 1.07 Acknowledgements

We would like to acknowledge Dhylan Patel, Antoine Lain and Nazinin Faghih-Mirzaei from the Posma group in the Division of Systems Medicine, Imperial College for developing the microbiome annotation pipeline, and for assistance with implementing it on our dataset.

## References

1. Owolade AJJ, Abdullateef RO, Adesola RO, Olaloye ED. Malnutrition: An underlying health condition faced in sub Saharan Africa: Challenges and recommendations. Ann Med Surg [Internet]. 2022 Oct [cited 2023 Aug 23];82. Available from: https://journals.lww.com/10.1016/j.amsu.2022.104769

2. Chege PM, Kimiywe JO, Ndungu ZW. Influence of culture on dietary practices of children under five years among Maasai pastoralists in Kajiado, Kenya. Int J Behav Nutr Phys Act. 2015 Dec;12(1):131.

3. Reddy S, Anitha M. Culture and its Influence on Nutrition and Oral Health. Biomed Pharmacol J. 2015 Oct 25;8(october Spl Edition):613–20.

4. Bain LE, Awah PK, Geraldine N, Kindong NP, Sigal Y, Bernard N, et al. Malnutrition in Sub-Saharan Africa: burden, causes and prospects. Pan Afr Med J [Internet]. 2013 [cited 2023 Aug 23];15. Available from: http://www.panafrican-med-journal.com/content/article/15/120/full/

5. Drammeh W, Hamid NA, Rohana AJ. Determinants of Household Food Insecurity and Its Association with Child Malnutrition in Sub-Saharan Africa: A Review of the Literature. Curr Res Nutr Food Sci J. 2019 Dec 31;7(3):610–23.

6. Black RE, Victora CG, Walker SP, Bhutta ZA, Christian P, de Onis M, et al. Maternal and child undernutrition and overweight in low-income and middle-income countries. The Lancet. 2013 Aug;382(9890):427–51.

7. Zyambo K, Hodges P, Chandwe K, Chisenga CC, Mayimbo S, Amadi B, et al. Selenium status in adults and children in Lusaka, Zambia. Heliyon. 2022 Jun;8(6):e09782.

8. Azeez TA. Obesity in Africa: The challenges of a rising epidemic in the midst of dwindling resources. Obes Med. 2022 May;31:100397.

9. De Vos WM, Tilg H, Van Hul M, Cani PD. Gut microbiome and health: mechanistic insights. Gut. 2022 May;71(5):1020–32.

10. Gomaa EZ. Human gut microbiota/microbiome in health and diseases: a review. Antonie Van Leeuwenhoek. 2020 Dec;113(12):2019–40.

11. Hou K, Wu ZX, Chen XY, Wang JQ, Zhang D, Xiao C, et al. Microbiota in health and diseases. Signal Transduct Target Ther. 2022 Apr 23;7(1):135.

12. Kortekangas E, Fan YM, Chaima D, Lehto KM, Malamba-Banda C, Matchado A, et al. Associations between Gut Microbiota and Intestinal Inflammation, Permeability and Damage in Young Malawian Children. J Trop Pediatr. 2022 Feb 3;68(2):fmac012.

13. Boicean A, Birlutiu V, Ichim C, Anderco P, Birsan S. Fecal Microbiota Transplantation in Inflammatory Bowel Disease. Biomedicines. 2023 Mar 27;11(4):1016.

14. Allegretti JR, Fischer M, Sagi SV, Bohm ME, Fadda HM, Ranmal SR, et al. Fecal Microbiota Transplantation Capsules with Targeted Colonic Versus Gastric Delivery in Recurrent Clostridium difficile Infection: A Comparative Cohort Analysis of High and Lose Dose. Dig Dis Sci. 2019 Jun 5;64(6):1672–8.

15. Aroniadis OC, Brandt LJ, Oneto C, Feuerstadt P, Sherman A, Wolkoff AW, et al. Faecal microbiota transplantation for diarrhoea-predominant irritable bowel syndrome: a double-blind, randomised, placebo-controlled trial. Lancet Gastroenterol Hepatol. 2019 Sep 1;4(9):675–85.

16. Gehrig JL, Venkatesh S, Chang HW, Hibberd MC, Kung VL, Cheng J, et al. Effects of microbiota-directed foods in gnotobiotic animals and undernourished children. Science. 2019 Jul 12;365(6449):eaau4732.

17. Chen RY, Mostafa I, Hibberd MC, Das S, Mahfuz M, Naila NN, et al. A Microbiota-Directed Food Intervention for Undernourished Children. N Engl J Med. 2021 Apr 22;384(16):1517–28.

18. Blanton LV, Barratt MJ, Charbonneau MR, Ahmed T, Gordon JI. Childhood undernutrition, the gut microbiota, and microbiota-directed therapeutics. Science. 2016 Jun 24;352(6293):1533–1533.

19. Kane AV, Dinh DM, Ward HD. Childhood malnutrition and the intestinal microbiome. Pediatr Res. 2015 Jan;77(1–2):256–62.

20. Dje Kouadio DK, Wieringa F, Greffeuille V, Humblot C. Bacteria from the gut influence the host micronutrient status. Crit Rev Food Sci Nutr. 2023 Jun 27;1–16.

21. Barratt MJ, Ahmed T, Gordon JI. Gut microbiome development and childhood undernutrition. Cell Host Microbe. 2022 May;30(5):617–26.

22. Iddrisu I, Monteagudo-Mera A, Poveda C, Pyle S, Shahzad M, Andrews S, et al. Malnutrition and gut microbiota in children. Nutrients. 2021 Aug 1;13(8).

23. Kennedy EA, Holtz LR. Gut virome in early life: origins and implications. Curr Opin Virol. 2022 Aug;55:101233.

24. Fontaine F, Turjeman S, Callens K, Koren O. The intersection of undernutrition, microbiome, and child development in the first years of life. Nat Commun. 2023 Jun 15;14(1):3554.

25. de Clercq NC, Groen AK, Romijn JA, Nieuwdorp M. Gut Microbiota in Obesity and Undernutrition. Adv Nutr Bethesda Md. 2016 Nov;7(6):1080–9.

26. Million M, Diallo A, Raoult D. Gut microbiota and malnutrition. Microb Pathog. 2017 May;106:127–38.

27. McGuire MK, McGuire MA. Microbiomes and Childhood Malnutrition: What Is the Evidence? Ann Nutr Metab. 2021;77(Suppl. 3):36–48.

28. Jones HJ, Bourke CD, Swann JR, Robertson RC. Malnourished Microbes: Host–Microbiome Interactions in Child Undernutrition. Annu Rev Nutr. 2023 Aug 21;43(1):327–53.

29. Borman AM, Johnson EM. Name Changes for Fungi of Medical Importance, 2020 to 2021. Humphries RM, editor. J Clin Microbiol. 2023 Jun 20;61(6):e00330–22.

30. Prinzi AM, Moore NM. Change of Plans: Overview of Bacterial Taxonomy, Recent Changes of Medical Importance, and Potential Areas of Impact. Open Forum Infect Dis. 2023 Jul;10(7):ofad269.

31. Relich RF, Loeffelholz MJ. Taxonomic Changes for Human Viruses, 2020 to 2022. Humphries RM, editor. J Clin Microbiol. 2023 Jan 26;61(1):e00337–22.

32. Janda JM. Taxonomic update on proposed nomenclature and classification changes for bacteria of medical importance, 2015. Diagn Microbiol Infect Dis. 2016 Oct;86(2):123–7.

33. Munson E, Carroll KC. An Update on the Novel Genera and Species and Revised Taxonomic Status of Bacterial Organisms Described in 2016 and 2017. Kraft CS, editor. J Clin Microbiol. 2019 Feb;57(2):e01181–18.

34. Goodfellow I, Bengio Y, Courville A. Deep Learning. Cambridge, MA, USA: MIT Press; 2016.

35. Karkera N, Acharya S, Palaniappan SK. Leveraging pre-trained language models for mining microbiome-disease relationships. BMC Bioinformatics. 2023 Jul 19;24(1):290.

36. Chibuye M, Mende DR, Spijker R, Simuyandi M, Luchen CC, Bosomprah S, et al. Systematic review of associations between gut microbiome composition and stunting in under-five children. Npj Biofilms Microbiomes. 2024 May 23;10(1):1–13.

37. World Bank Country and Lending Groups – World Bank Data Help Desk [Internet]. [cited 2025 May 6]. Available from: https://datahelpdesk.worldbank.org/knowledgebase/articles/906519-world-bank-country-and-lending-groups

38. Islamaj R, Kwon D, Kim S, Lu Z. TeamTat: a collaborative text annotation tool. Nucleic Acids Res. 2020 Jul 2;48(W1):W5–11.

39. Patel D. icprofsensei/New_Pipeline_2023: Bug fixes [Internet]. Zenodo; 2023 [cited 2024 Apr 3]. Available from: https://zenodo.org/records/8223801

40. Beck T, Shorter T, Hu Y, Li Z, Sun S, Popovici CM, et al. Auto-CORPus: A Natural Language Processing Tool for Standardizing and Reusing Biomedical Literature. Front Digit Health. 2022 Feb 15;4:788124.

41. R Core Team. R: A Language and Environment for statistical Computing [Internet]. R Foundation for Statistical Computing. 2022. Available from: https://www.R-project.org/

42. Calder N, Walsh K, Olupot-Olupot P, Ssenyondo T, Muhindo R, Mpoya A, et al. Modifying gut integrity and microbiome in children with severe acute malnutrition using legume-based feeds (MIMBLE): A pilot trial. Cell Rep Med. 2021 May 18;2(5):100280.

43. Collard JM, Andrianonimiadana L, Habib A, Rakotondrainipiana M, Andriantsalama P, Randriamparany R, et al. High prevalence of small intestine bacteria overgrowth and asymptomatic carriage of enteric pathogens in stunted children in Antananarivo, Madagascar. Azman AS, editor. PLoS Negl Trop Dis. 2022 May 9;16(5):e0009849.

44. Desai C, Handley SA, Rodgers R, Rodriguez C, Ordiz MI, Manary MJ, et al. Growth velocity in children with Environmental Enteric Dysfunction is associated with specific bacterial and viral taxa of the gastrointestinal tract in Malawian children. Melby PC, editor. PLoS Negl Trop Dis. 2020 Jun 23;14(6):e0008387.

45. Ismail NA, Ragab SH, ElBaky AA, Shoeib ARS, Alhosary Y, Fekry D. Frequency of Firmicutes and Bacteroidetes in gut microbiota in obese and normal weight Egyptian children and adults. Arch Med Sci. 2011;3:501–7.

46. Ordiz MI, May TD, Mihindukulasuriya K, Martin J, Crowley J, Tarr PI, et al. The effect of dietary resistant starch type 2 on the microbiota and markers of gut inflammation in rural Malawi children. Microbiome. 2015 Dec 3;3(1):37.

47. Ordiz MI, Stephenson K, Agapova S, Wylie KM, Maleta K, Martin J, et al. Environmental Enteric Dysfunction and the Fecal Microbiota in Malawian Children. Am J Trop Med Hyg. 2017 Feb 8;96(2):473–6.

48. Vonaesch P, Morien E, Andrianonimiadana L, Sanke H, Mbecko JR, Huus KE, et al. Stunted childhood growth is associated with decompartmentalization of the gastrointestinal tract and overgrowth of oropharyngeal taxa. Proc Natl Acad Sci U S A. 2018 Sep 4;115(36):E8489–98.

49. Vonaesch P, Araújo JR, Gody JC, Mbecko JR, Sanke H, Andrianonimiadana L, et al. Stunted children display ectopic small intestinal colonization by oral bacteria, which cause lipid malabsorption in experimental models. Proc Natl Acad Sci U S A. 2022 Oct 11;119(41):e2209589119.

50. Mweetwa MN. Africa-MetaAnalysis Code [Internet]. 2024 [cited 2024 Jul 22]. Available from: https://github.com/Monica-Mweetwa/Africa-MetaAnalysis-Archive.git

51. Paradis E, Schliep K. ape 5.0: an environment for modern phylogenetics and evolutionary analyses in R. Schwartz R, editor. Bioinformatics. 2019 Feb 1;35(3):526–8.

52. Yu G. Data Integration, Manipulation and Visualization of Phylogenetic Trees [Internet]. 1st ed. Boca Raton: Chapman and Hall/CRC; 2022 [cited 2024 Feb 1]. Available from: https://www.taylorfrancis.com/books/9781003279242

53. Scott sherrill-Mix. taxonomizr: Functions to Work with NCBI Accessions and Taxonomy [Internet]. 2023. Available from: https://CRAB.R-project.rog/package=taxonomizr

54. Hadley Wickham. ggplot2:Elegant Graphics for Data Analysis [Internet]. Springer-Verlag New York; 2016. Available from: https://ggplot2.tidyverse.org

55. Sterne JAC, Savović J, Page MJ, Elbers RG, Blencowe NS, Boutron I, et al. RoB 2: a revised tool for assessing risk of bias in randomised trials. BMJ. 2019 Aug 28;l4898.

56. Higgins JPT, Morgan RL, Rooney AA, Taylor KW, Thayer KA, Silva RA, et al. A tool to assess risk of bias in non-randomized follow-up studies of exposure effects (ROBINS-E). Environ Int. 2024 Apr;186:108602.

57. Sterne JA, Hernán MA, Reeves BC, Savović J, Berkman ND, Viswanathan M, et al. ROBINS-I: a tool for assessing risk of bias in non-randomised studies of interventions. BMJ. 2016 Oct 12;i4919.

58. GitHub [Internet]. [cited 2024 Oct 30]. Python. Available from: https://github.com/python

59. Popovic A, Bourdon C, Wang PW, Guttman DS, Soofi S, Bhutta ZA, et al. Micronutrient supplements can promote disruptive protozoan and fungal communities in the developing infant gut. Nat Commun. 2021 Nov 18;12(1):6729.

60. WHO. Levels and trends in child malnutrition: UNICEF/WHO/The World Bank Group joint child malnutrition estimates: key findings of the 2021 edition [Internet]. 2021. Report No.: ISBN: 9789240025257. Available from: https://www.who.int/publications/i/item/9789240025257

61. Anders JL, Moustafa MAM, Mohamed WMA, Hayakawa T, Nakao R, Koizumi I. Comparing the gut microbiome along the gastrointestinal tract of three sympatric species of wild rodents. Sci Rep. 2021 Oct 7;11(1):19929.

62. Mulenga C, Sviben S, Chandwe K, Amadi B, Kayamba V, Fitzpatrick JAJ, et al. Epithelial Abnormalities in the Small Intestine of Zambian Children With Stunting. Front Med. 2022 Mar 16;9:849677.

63. Nejati S, Wang J, Sedaghat S, Balog NK, Long AM, Rivera UH, et al. Smart capsule for targeted proximal colon microbiome sampling. Acta Biomater. 2022 Dec;154:83–96.

64. Waimin JF, Nejati S, Jiang H, Qiu J, Wang J, Verma MS, et al. Smart capsule for non-invasive sampling and studying of the gastrointestinal microbiome. RSC Adv. 2020;10(28):16313–22.

65. Shalon D, Culver RN, Grembi JA, Folz J, Treit PV, Shi H, et al. Profiling the human intestinal environment under physiological conditions. Nature. 2023 May 18;617(7961):581–91.

66. Rhoads A, Au KF. PacBio Sequencing and Its Applications. Genomics Proteomics Bioinformatics. 2015 Oct;13(5):278–89.

67. Jain M, Olsen HE, Paten B, Akeson M. The Oxford Nanopore MinION: delivery of nanopore sequencing to the genomics community. Genome Biol. 2016 Dec;17(1):239.

68. Chetty A, Blekhman R. Multi-omic approaches for host-microbiome data integration. Gut Microbes. 2024;16(1):2297860.

69. Jiang D, Armour CR, Hu C, Mei M, Tian C, Sharpton TJ, et al. Microbiome Multi-Omics Network Analysis: Statistical Considerations, Limitations, and Opportunities. Front Genet. 2019 Nov 8;10:995.

70. Heyworth B, Brown J. Jejunal microflora in malnourished Gambian children. Arch Dis Child. 1975 Jan 1;50(1):27–33.

71. Omoike IU, Abiodun PO. Upper Small Intestinal Microflora in Diarrhea and Malnutrition in Nigerian Children: J Pediatr Gastroenterol Nutr. 1989 Oct;9(3):314–21.

72. Armstrong G, Rahman G, Martino C, McDonald D, Gonzalez A, Mishne G, et al. Applications and Comparison of Dimensionality Reduction Methods for Microbiome Data. Front Bioinforma. 2022 Feb 24;2:821861.

73. Reyes A, Blanton LV, Cao S, Zhao G, Manary M, Trehan I, et al. Gut DNA viromes of Malawian twins discordant for severe acute malnutrition. Proc Natl Acad Sci U S A. 2015 Sep 22;112(38):11941–6.

74. Alou MT, Rathored J, Michelle C, Dubourg G, Andrieu C, Armstrong N, et al. Inediibacterium massiliense gen. nov., sp. nov., a new bacterial species isolated from the gut microbiota of a severely malnourished infant. Antonie Van Leeuwenhoek. 2017 Jun;110(6):737–50.

75. Pham TPT, Cadoret F, Alou MT, Brah S, Diallo BA, Diallo A, et al. “Urmitella timonensis” gen. nov., sp. nov., “Blautia marasmi” sp. nov., “Lachnoclostridium pacaense” sp. nov., “Bacillus marasmi” sp. nov. and “Anaerotruncus rubiinfantis” sp. nov., isolated from stool samples of undernourished African children. New Microbes New Infect. 2017 May;17:84–8.

76. Smith MI, Yatsunenko T, Manary MJ, Trehan I, Mkakosya R, Cheng J, et al. Gut Microbiomes of Malawian Twin Pairs Discordant for Kwashiorkor. Science. 2013 Feb 1;339(6119):548–54.

77. Tidjani Alou M, Rathored J, Lagier JC, Khelaifia S, Michelle C, Sokhna C, et al. Rubeoparvulum massiliense gen. nov., sp. nov., a new bacterial genus isolated from the human gut of a Senegalese infant with severe acute malnutrition. New Microbes New Infect. 2017 Jan;15:49–60.

78. Laursen MF, Sakanaka M, von Burg N, Mörbe U, Andersen D, Moll JM, et al. Bifidobacterium species associated with breastfeeding produce aromatic lactic acids in the infant gut. Nat Microbiol. 2021 Nov;6(11):1367–82.

79. Sadeghpour Heravi F, Hu H. Bifidobacterium: Host-Microbiome Interaction and Mechanism of Action in Preventing Common Gut-Microbiota-Associated Complications in Preterm Infants: A Narrative Review. Nutrients. 2023 Jan 30;15(3):709.

80. Schwartz DJ, Langdon A, Sun X, Langendorf C, Berthé F, Grais RF, et al. Effect of amoxicillin on the gut microbiome of children with severe acute malnutrition in Madarounfa, Niger: a retrospective metagenomic analysis of a placebo-controlled trial. Lancet Microbe. 2023 Nov;4(11):e931–42.

81. Perin J, Burrowes V, Almeida M, Ahmed S, Haque R, Parvin T, et al. A Retrospective Case-Control Study of the Relationship between the Gut Microbiota, Enteropathy, and Child Growth. Am J Trop Med Hyg. 2020 Jul;103(1):520–7.

82. Balaji V, Dinh DM, Kane AV, Soofi S, Ahmed I, Rizvi A, et al. Longitudinal Analysis of the Intestinal Microbiota among a Cohort of Children in Rural and Urban Areas of Pakistan. Nutrients. 2023 Feb 28;15(5):1213.

83. Shivakumar N, Sivadas A, Devi S, Jahoor F, McLaughlin J, Smith CP, et al. Gut microbiota profiles of young South Indian children: Child sex-specific relations with growth. PloS One. 2021;16(5):e0251803.

84. Balasubramaniam C, Mallappa RH, Singh DK, Chaudhary P, Bharti B, Muniyappa SK, et al. Gut bacterial profile in Indian children of varying nutritional status: a comparative pilot study. Eur J Nutr. 2021 Oct;60(7):3971–85.

85. Barratt MJ, Nuzhat S, Ahsan K, Frese SA, Arzamasov AA, Sarker SA, et al. Bifidobacterium infantis treatment promotes weight gain in Bangladeshi infants with severe acute malnutrition. Sci Transl Med. 2022 Apr 13;14(640):eabk1107.

86. Ghosh TS, Gupta SS, Bhattacharya T, Yadav D, Barik A, Chowdhury A, et al. Gut microbiomes of Indian children of varying nutritional status. PloS One. 2014;9(4):e95547.

87. Ottman N, Smidt H, de Vos WM, Belzer C. The function of our microbiota: who is out there and what do they do? Front Cell Infect Microbiol. 2012;2:104.

88. Bressuire-Isoard C, Broussolle V, Carlin F. Sporulation environment influences spore properties in Bacillus: evidence and insights on underlying molecular and physiological mechanisms. FEMS Microbiol Rev. 2018 Sep 1;42(5):614–26.

89. Nicholson WL, Munakata N, Horneck G, Melosh HJ, Setlow P. Resistance of Bacillus Endospores to Extreme Terrestrial and Extraterrestrial Environments. Microbiol Mol Biol Rev. 2000 Sep;64(3):548–72.

90. Nabwera HM, Espinoza JL, Worwui A, Betts M, Okoi C, Sesay AK, et al. Interactions between fecal gut microbiome, enteric pathogens, and energy regulating hormones among acutely malnourished rural Gambian children. EBioMedicine. 2021 Nov 1;73.

91. Castro-Mejía JL, O’Ferrall S, Krych Ł, O’Mahony E, Namusoke H, Lanyero B, et al. Restitution of gut microbiota in Ugandan children administered with probiotics ( *Lactobacillus rhamnosus* GG and *Bifidobacterium animalis* subsp. *lactis* BB-12) during treatment for severe acute malnutrition. Gut Microbes. 2020 Jul 3;11(4):855–67.

92. Zambrana LE, McKeen S, Ibrahim H, Zarei I, Borresen EC, Doumbia L, et al. Rice bran supplementation modulates growth, microbiota and metabolome in weaning infants: a clinical trial in Nicaragua and Mali. Sci Rep. 2019 Dec 26;9(1):13919.

93. Aziz I, Noreen Z, Ijaz UZ, Gundogdu O, Hamid MH, Muhammad N, et al. A prospective study on linking diarrheagenic E. coli with stunted childhood growth in relation to gut microbiome. Sci Rep. 2023 Apr 26;13(1):6802.

94. Mostafa I, Hibberd MC, Hartman SJ, Hafizur Rahman MH, Mahfuz M, Hasan SMT, et al. A microbiota-directed complementary food intervention in 12-18-month-old Bangladeshi children improves linear growth. EBioMedicine. 2024 Jun;104:105166.

95. Camara A, Konate S, Tidjani Alou M, Kodio A, Togo AH, Cortaredona S, et al. Clinical evidence of the role of Methanobrevibacter smithii in severe acute malnutrition. Sci Rep. 2021 Dec 8;11(1):5426.

96. Donkor ES. Stroke in the 21 st Century: A Snapshot of the Burden, Epidemiology, and Quality of Life. Stroke Res Treat. 2018 Nov 27;2018:1–10.

97. Kristensen KHS, Wiese M, Rytter MJH, Özçam M, Hansen LH, Namusoke H, et al. Gut Microbiota in Children Hospitalized with Oedematous and Non-Oedematous Severe Acute Malnutrition in Uganda. PLoS Negl Trop Dis. 2016 Jan 15;10(1).

98. Tidjani Alou M, Million M, Traore SI, Mouelhi D, Khelaifia S, Bachar D, et al. Gut Bacteria Missing in Severe Acute Malnutrition, Can We Identify Potential Probiotics by Culturomics? Front Microbiol. 2017 May 23;8:899.

99. Monira S, Nakamura S, Gotoh K, Izutsu K, Watanabe H, Alam NH, et al. Gut Microbiota of Healthy and Malnourished Children in Bangladesh. Front Microbiol [Internet]. 2011 [cited 2024 Feb 9];2. Available from: http://journal.frontiersin.org/article/10.3389/fmicb.2011.00228/abstract

100. Caruso R, Lo BC, Núñez G. Host-microbiota interactions in inflammatory bowel disease. Nat Rev Immunol. 2020 Jul;20(7):411–26.

101. Zambruni M, Ochoa TJ, Somasunderam A, Cabada MM, Morales ML, Mitreva M, et al. Stunting Is Preceded by Intestinal Mucosal Damage and Microbiome Changes and Is Associated with Systemic Inflammation in a Cohort of Peruvian Infants. Am J Trop Med Hyg. 2019 Nov;101(5):1009–17.

102. Lo Presti A, Del Chierico F, Altomare A, Zorzi F, Monteleone G, Putignani L, et al. Phylogenetic analysis of *Prevotella copri* from fecal and mucosal microbiota of IBS and IBD patients. Ther Adv Gastroenterol. 2023 Jan;16:175628482211363.

103. Dinh DM, Ramadass B, Kattula D, Sarkar R, Braunstein P, Tai A, et al. Longitudinal Analysis of the Intestinal Microbiota in Persistently Stunted Young Children in South India. Parkinson J, editor. PLOS ONE. 2016 May 26;11(5):e0155405.

104. Liu J, Zhou L, Sun L, Ye X, Ma M, Dou M, et al. Association Between Intestinal Prevotella copri Abundance and Glycemic Fluctuation in Patients with Brittle Diabetes. Diabetes Metab Syndr Obes. 2023 Jun;Volume 16:1613–21.

105. Robertson RC, Edens TJ, Carr L, Mutasa K, Gough EK, Evans C, et al. The gut microbiome and early-life growth in a population with high prevalence of stunting. Nat Commun. 2023 Feb 14;14(1):654.

106. Doumatey AP, Adeyemo A, Zhou J, Lei L, Adebamowo SN, Adebamowo C, et al. Gut Microbiome Profiles Are Associated With Type 2 Diabetes in Urban Africans. Front Cell Infect Microbiol. 2020 Feb 25;10.

107. Könönen E, Gursoy UK. Oral Prevotella Species and Their Connection to Events of Clinical Relevance in Gastrointestinal and Respiratory Tracts. Front Microbiol. 2022 Jan 6;12:798763.

108. Balakrishnan B, Luckey D, Bodhke R, Chen J, Marietta E, Jeraldo P, et al. Prevotella histicola Protects From Arthritis by Expansion of Allobaculum and Augmenting Butyrate Production in Humanized Mice. Front Immunol. 2021 May 4;12:609644.

109. Balakrishnan B, Luckey D, Taneja V. Autoimmunity-Associated Gut Commensals Modulate Gut Permeability and Immunity in Humanized Mice. Mil Med. 2019 Mar 1;184(Suppl 1):529–36.

110. de Goffau MC, Jallow AT, Sanyang C, Prentice AM, Meagher N, Price DJ, et al. Gut microbiomes from Gambian infants reveal the development of a non-industrialized Prevotella-based trophic network. Nat Microbiol. 2022 Jan 31;7(1):132–44.

111. Oduaran OH, Tamburini FB, Sahibdeen V, Brewster R, Gómez-Olivé FX, Kahn K, et al. Gut microbiome profiling of a rural and urban South African cohort reveals biomarkers of a population in lifestyle transition. BMC Microbiol. 2020 Dec 31;20(1):330.

112. Tett A, Huang KD, Asnicar F, Fehlner-Peach H, Pasolli E, Karcher N, et al. The Prevotella copri Complex Comprises Four Distinct Clades Underrepresented in Westernized Populations. Cell Host Microbe. 2019 Nov 13;26(5):666–679.e7.

113. Robertson RC, Manges AR, Finlay BB, Prendergast AJ. The Human Microbiome and Child Growth – First 1000 Days and Beyond. Trends Microbiol. 2019 Feb;27(2):131–47.

114. Geurtsen J, de Been M, Weerdenburg E, Zomer A, McNally A, Poolman J. Genomics and pathotypes of the many faces of *Escherichia coli*. FEMS Microbiol Rev. 2022 Nov 2;46(6):fuac031.

115. Amadi B, Zyambo K, Chandwe K, Besa E, Mulenga C, Mwakamui S, et al. Adaptation of the small intestine to microbial enteropathogens in Zambian children with stunting. Nat Microbiol. 2021 Feb 15;6(4):445–54.

116. Gazi MdA, Alam MdA, Fahim SM, Wahid BZ, Khan SS, Islam MdO, et al. Infection With Escherichia Coli Pathotypes Is Associated With Biomarkers of Gut Enteropathy and Nutritional Status Among Malnourished Children in Bangladesh. Front Cell Infect Microbiol. 2022 Jul 6;12:901324.

117. Onubi OJ, Poobalan AS, Dineen B, Marais D, McNeill G. Effects of probiotics on child growth: a systematic review. J Health Popul Nutr. 2015 Dec;34(1):8.

118. Mayneris-Perxachs J, Cardellini M, Hoyles L, Latorre J, Davato F, Moreno-Navarrete JM, et al. Iron status influences non-alcoholic fatty liver disease in obesity through the gut microbiome. Microbiome. 2021 May 7;9(1):104.

119. Mei H, Li N, Zhang Y, Zhang D, Peng AN, Tan YF, et al. Gut Microbiota Diversity and Overweight/Obesity in Infancy: Results from a Nested Case-control Study. Curr Med Sci. 2022 Feb;42(1):210–6.

120. Stefura T, Zapała B, Gosiewski T, Skomarovska O, Dudek A, Pędziwiatr M, et al. Differences in Compositions of Oral and Fecal Microbiota between Patients with Obesity and Controls. Med Kaunas Lith. 2021 Jun 30;57(7):678.

121. Holmberg SM, Feeney RH, Prasoodanan P.K. V, Puértolas-Balint F, Singh DK, Wongkuna S, et al. The gut commensal Blautia maintains colonic mucus function under low-fiber consumption through secretion of short-chain fatty acids. Nat Commun. 2024 Apr 25;15(1):3502.

122. Liu X, Mao B, Gu J, Wu J, Cui S, Wang G, et al. Blautia-a new functional genus with potential probiotic properties? Gut Microbes. 2021;13(1):1–21.

123. Faden H. The Role of Faecalibacterium, Roseburia, and Butyrate in Inflammatory Bowel Disease. Dig Dis. 2022;40(6):793–5.

124. Nie K, Ma K, Luo W, Shen Z, Yang Z, Xiao M, et al. Roseburia intestinalis: A Beneficial Gut Organism From the Discoveries in Genus and Species. Front Cell Infect Microbiol. 2021;11:757718.

125. Tamanai-Shacoori Z, Smida I, Bousarghin L, Loreal O, Meuric V, Fong SB, et al. Roseburia spp.: a marker of health? Future Microbiol. 2017 Feb;12:157–70.

126. Oren A, Arahal DR, Rosselló-Móra R, Sutcliffe IC, Moore ERB. Preparing a revision of the International Code of Nomenclature of Prokaryotes. Int J Syst Evol Microbiol [Internet]. 2021 Jan 1 [cited 2024 Feb 9];71(1). Available from: https://www.microbiologyresearch.org/content/journal/ijsem/10.1099/ijsem.0.004598

127. Mirzayi C, Renson A, Zohra F, Elsafoury S, Geistlinger L, Kasselman LJ, et al. Reporting guidelines for human microbiome research: the STORMS checklist. Nat Med. 2021 Nov;27(11):1885–92.

128. Frontiers | Reducing bias in microbiome research: Comparing methods from sample collection to sequencing [Internet]. [cited 2025 May 8]. Available from: https://www.frontiersin.org/journals/microbiology/articles/10.3389/fmicb.2023.1094800/full

