## Supplementary material for "A META-ANALYSIS OF GUT MICROBIOME RESEARCH IN MALNOURISHED AFRICAN POPULATIONS: A NATURAL LANGUAGE PROCESSING APPROACH": S1_Data

### **A narrative review of gut microbiome research in malnourished Sub-Saharan African populations using natural language processing meta-analysis approach**

### Mweetwa et al

### Supplementary Methodology and Figures

**Detailed search strategy employed in PubMed:**

1. Sub-Saharan Africa

(gut microflora OR gut microbiota OR gut microbiome OR gut microbiomics OR gut metagenome OR gut metagenomics OR gut resistome OR gut phageome OR gut viromics OR gut virome OR gut mycobiome OR gut metatranscriptomics) AND (Angola OR Burundi OR Benin “Burkina Faso” OR “Central African Republic” OR Côte d’Ivoire OR "Ivory Coast" OR Cameroon OR Chad OR Congo OR "Democratic Republic of the Congo" OR Zaire OR “The Republic of Congo” OR Comoros OR "Iles Comores" OR "Comoro Islands" OR Cabo Verde OR OR “Cape Verde” OR “República de Cabo Verde” OR Eswatini OR Swaziland OR Eritrea OR Ethiopia OR Ghana OR Guinea OR “The Gambia” OR Gambia OR “Guinea-Bissau” OR Kenya OR Liberia OR Lesotho OR Madagascar OR Mali OR Mozambique OR Mauritania OR Malawi OR Niger OR Nigeria OR Rwanda OR Sudan OR “South Sudan” OR Senegal OR “Sierra Leone” OR Somalia OR “São Tomé and Príncipe” OR Togo OR Tanzania OR Uganda OR Zambia OR Zimbabwe) AND (humans OR human) AND (Malnutrition OR Stunting OR Marasmus OR Kwashiorkor OR Wasting OR Undernourished OR Undernutrition OR Underweight OR “Vitamin Deficiency” OR Obesity OR Overweight).

To understand if microbial features are region specific, additional searches for LMICs outside Africa was done as follows:

2. Europe & Central Asia

(gut microflora OR gut microbiota OR gut microbiome OR gut microbiomics OR gut metagenome OR gut metagenomics OR gut resistome OR gut phageome OR gut viromics OR gut virome OR gut mycobiome OR gut metatranscriptomics) AND (“Kyrgyz Republic” OR Kyrgyzstan OR Tajikistan OR Ukraine OR Uzbekistan) AND (humans OR human) AND (Malnutrition OR Stunting OR Marasmus OR Kwashiokor OR Wasting OR Undernourished OR Undernutrition OR Underweight OR “Vitamin Deficiency” OR Obesity OR Overweight)

2. East Asia & Pacific

(gut microflora OR gut microbiota OR gut microbiome OR gut microbiomics OR gut metagenome OR gut metagenomics OR gut resistome OR gut phageome OR gut viromics OR gut virome OR gut mycobiome OR gut metatranscriptomics) AND (Micronesia OR “Federated States of Micronesia” OR Cambodia OR Kiribati OR Laos OR “Lao People's Democratic Republic” OR lao OR Myanmar OR Mongolia OR Philippines OR “Papua New Guinea” OR “North Korea” OR “Democratic People's Republic of Korea” OR “Solomon Islands” OR “East Timor” OR “Timor-Leste” OR “Timor Leste” OR Vietnam OR Vanuatu OR Samoa) AND (humans OR human) AND (Malnutrition OR Stunting OR Marasmus OR Kwashiokor OR Wasting OR Undernourished OR Undernutrition OR Underweight OR “Vitamin Deficiency” OR Obesity OR Overweight)

3. South Asia

(gut microflora OR gut microbiota OR gut microbiome OR gut microbiomics OR gut metagenome OR gut metagenomics OR gut resistome OR gut phageome OR gut viromics OR gut virome OR gut mycobiome OR gut metatranscriptomics) AND (Afghanistan OR Bangladesh OR Bhutan OR India OR “Sri Lanka” OR Nepal OR Pakistan) AND (humans OR human) AND (Malnutrition OR Stunting OR Marasmus OR Kwashiokor OR Wasting OR Undernourished OR Undernutrition OR Underweight OR “Vitamin Deficiency” OR Obesity OR Overweight)

4. Latin America & Caribbean

(gut microflora OR gut microbiota OR gut microbiome OR gut microbiomics OR gut metagenome OR gut metagenomics OR gut resistome OR gut phageome OR gut viromics OR gut virome OR gut mycobiome OR gut metatranscriptomics) AND (Bolivia OR Honduras OR Haiti OR “Republic of Haiti” OR Nicaragua) AND (humans OR human) AND (Malnutrition OR Stunting OR Marasmus OR Kwashiorkor OR Wasting OR Undernourished OR Undernutrition OR Underweight OR “Vitamin Deficiency” OR Obesity OR Overweight)

5. Middle East & North Africa

(gut microflora OR gut microbiota OR gut microbiome OR gut microbiomics OR gut metagenome OR gut metagenomics OR gut resistome OR gut phageome OR gut viromics OR gut virome OR gut mycobiome OR gut metatranscriptomics) AND (Djibouti OR Algeria OR Egypt OR Iran OR Jordan OR Lebanon OR Morocco OR “Syrian Arab Republic” Or Syria OR Tunisia OR Yemen OR “Republic of Yemen”) AND (humans OR human) AND (Malnutrition OR Stunting OR Marasmus OR Kwashiorkor OR Wasting OR Undernourished OR Undernutrition OR Underweight OR “Vitamin Deficiency” OR Obesity OR Overweight).

### Supplementary figures

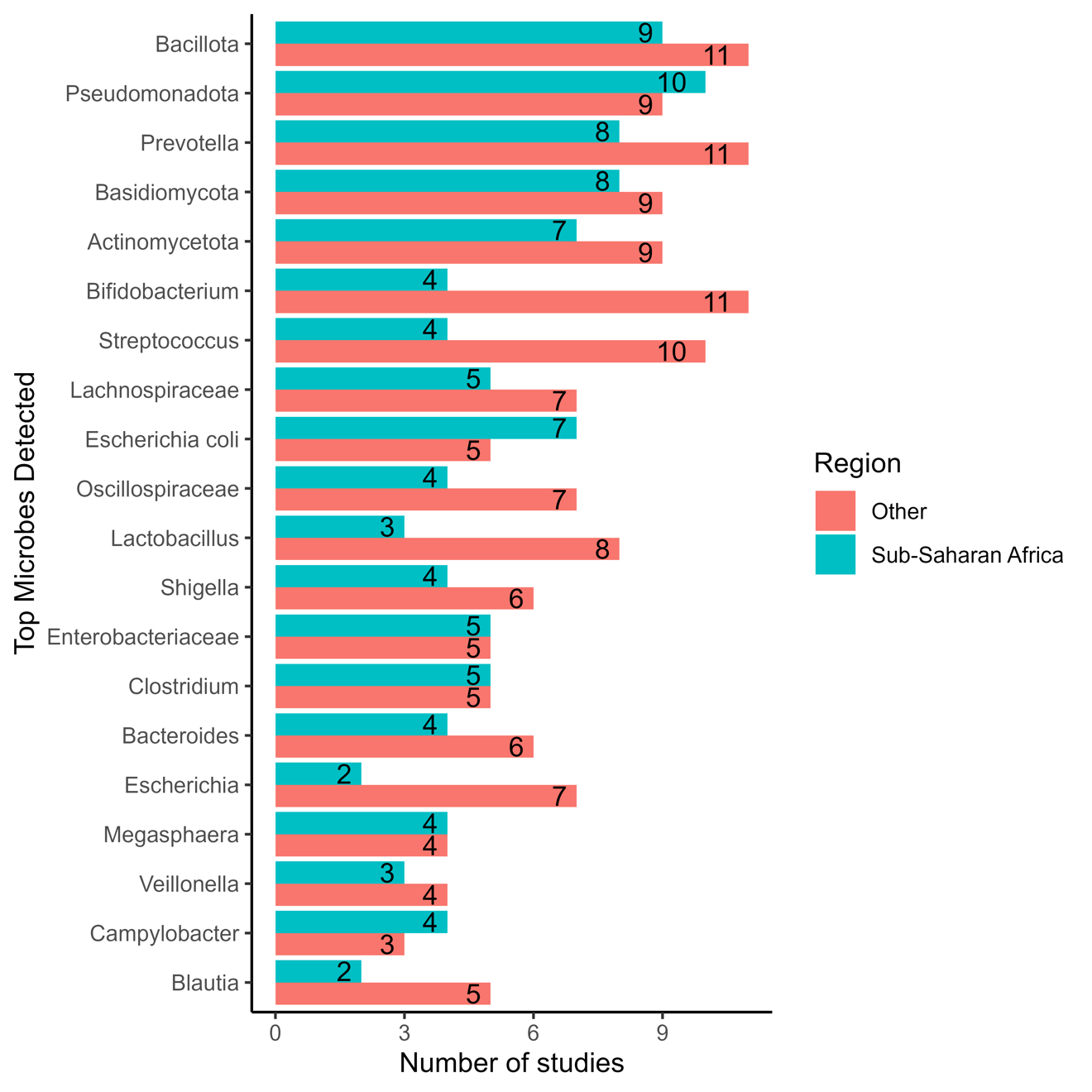

Supplementary figure 1: the distribution of the most frequently reported microbes.
